# Hydrodynamics of transient cell-cell contact: The role of membrane permeability and active protrusion length

**DOI:** 10.1101/367987

**Authors:** Kai Liu, Brian Chu, Jay Newby, Elizabeth L. Read, John Lowengrub, Jun Allard

## Abstract

In many biological settings, two or more cells come into physical contact to form a cell-cell interface. In some cases, the cell-cell contact must be transient, forming on timescales of seconds. One example is offered by the T cell, an immune cell which must attach to the surface of other cells in order to decipher information about disease. The aspect ratio of these interfaces (tens of nanometers thick and tens of micrometers in diameter) puts them into the thin-layer limit, or “lubrication limit”, of fluid dynamics. A key question is how the receptors and ligands on opposing cells come into contact. What are the relative roles of thermal undulations of the plasma membrane and deterministic forces from active filopodia? We use a computational fluid dynamics algorithm capable of simulating 10-nanometer-scale fluid-structure interactions with thermal fluctuations up to seconds-and microns-scales. We use this to simulate two opposing membranes, variously including thermal fluctuations, active forces, and membrane permeability. In some regimes dominated by thermal fluctuations, proximity is a rare event, which we capture by computing mean first-passage times using a Weighted Ensemble rare-event computational method. Our results demonstrate that the time-to-contact increases for smaller cell-cell distances (where the thin-layer effect is strongest), leading to an optimal initial cell-cell separation for fastest receptor-ligand binding. We reproduce a previous experimental observation that fluctuation spatial scales are largely unaffected, but timescales are dramatically slowed, by the thin-layer effect. We also find that membrane permeability would need to be above physiological levels to abrogate the thin-layer effect.

**Author summary:** The elastohydrodynamics of water in and around cells is playing an increasingly recognized role in biology. In this work, we investigate the flow of extracellular fluid in between cells during the formation of a cell-cell contact, to determine whether its necessary evacuation as the cells approach is a rate-limiting step before molecules on either cell can interact. To overcome the computational challenges associated with simulating fluid in this mechanically soft, stochastic and high-aspect-ratio environment, we extend a computational framework where the cell plasma membranes are treated as immersed boundaries in the fluid, and combine this with computational methods for simulating stochastic rare events in which an ensemble of simulations are given weights according to their probability. We find that the internal dynamics of the membranes has speeds in approximately microseconds, but that as the cells approach, a new slow timescale of approximately milliseconds is introduced. Thermal undulations nor typical amounts of membrane permeability can overcome the timescale, but active forces, e.g., from the cytoskeleton, can. Our results suggest an explanation for differences in molecular interactions in live cells compared to in vitro reconstitution experiments.

## Introduction

In many biological processes, two or more cells come into physical contact to form a cell-cell interface. These include cell-cell contacts like those in the epithelium [1, 2] that change on timescales of hours, and also transient contacts that form on seconds timescales, including those formed by lymphocytes and other immune cells that must interrogate many cells rapidly [3, 4]. A fundamental question for all cell-cell interfaces is how receptors and ligands come into contact, despite being separated by extracellular fluid, various large surface molecules like ectodomains of membrane proteins, and other structures in the negatively-charged glycocalyx. The contribution of large surface molecules has received most attention, for example producing spatial pattern formation based on molecular size [5–9] of the T cell receptor (TCR) and the immunotherapy target PD-1 [10]. In this work, we focus on the role of the fluid [11–14].

To highlight the potential importance of the hydrodynamics of extracellular fluid at an interface, we perform a preliminary calculation (unrealistically) assuming cells are rigid, impermeable spheres of radius *r*_cell_. In order to bring these cells into close contact, a force *F* pushes them together, as shown in Fig. 1A. This fluid dynamics problem can be solved analytically for the separation distance *z*, yielding [15, 16]

**Fig 1.**
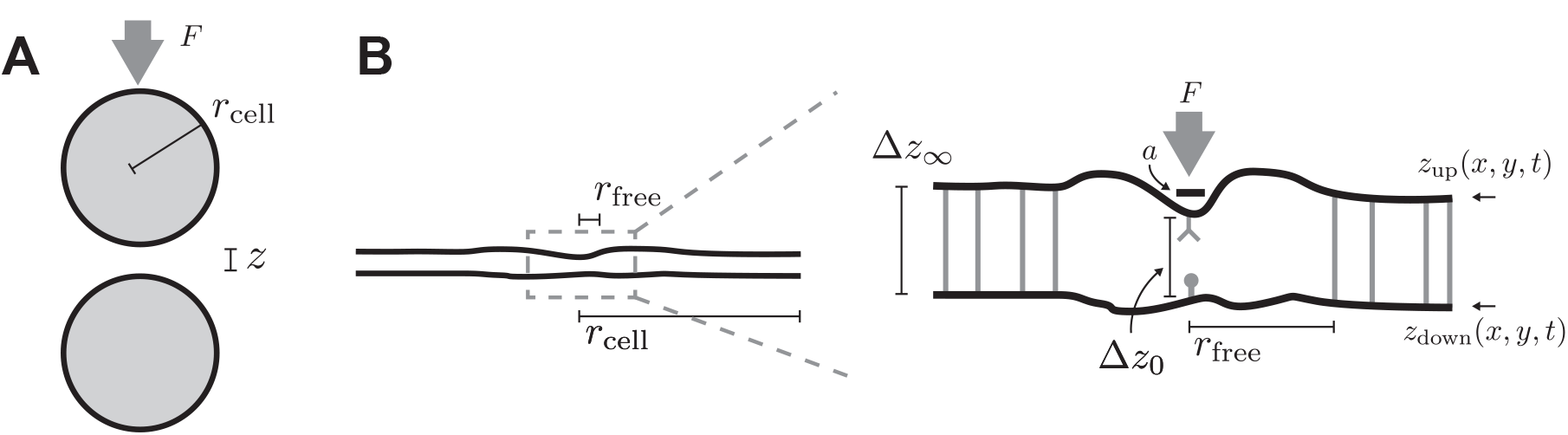
(A) Two cells, here depicted as spheres, pushed together by a force *F*. (B) Schematic of model geometry. Both cells have radius *r*_cell_ much larger than the cell-cell separation distance (left). We assume the cells are held apart by nonspecific adhesion molecules with size Δ*z*_∞_, which we refer to as the far-field separation. Near the receptor, there is a region free of nonspecific adhesion molecules of radius *r*_free_, which is related to the surface density of non-specific adhesion molecules 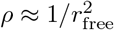. The membrane separation distance at the receptor is Δ*z*_0_. In simulations with active forces, the force *F* is applied to a circular area of the top membrane with radius *a*.

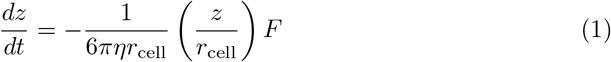
where *η* is the extracellular fluid viscosity. This equation is reminiscent of the Stokes drag formula for a sphere in free fluid, but modified by a factor (*z/r*_cell_) ∼ (10 *μ*m*/*10 nm) ∼ 10^3^. In other words, the force required to move two cells together is increased by a thousand-fold, a strikingly large correction. This observation, known as the “lubrication limit”, “confinement effect” or “thin-layer effect” [11, 15, 16], heuristically arises because a small change in *z* requires incompressible fluid to move a large distance to outside the interface.

The cell surface is not a rigid sphere, but a deformable membrane subject to thermal undulations, active forces, and hydraulic permeability due largely to membrane inclusions like aquaporins. In this context, we ask, what is the role of the fluid in close-contact formation? Are thermal undulations sufficient for receptor proximity? Are typical F-actin filopodial forces, ∼10 picoNewtons [17, 18], sufficient for receptor proximity? And how much force is required for rapid proximity (<1 second)? If there is a significant thin-layer effect, the force required will increase for smaller cell-cell distances, but larger distances require longer protrusions, suggesting the possibility of an optimal “attack range” which might explain the biological benefit of filopodia. If the membrane is permeable to extracellular fluid [19], how much permeability is required for rapid proximity? Factors that influence permeability, such as aquaporins, are under regulation [20], differentially localized, and impact cell processes including cancer angiogenesis [21], raising the possibility that cell-cell contact can be regulated in this way.

In contrast to previous theoretical studies of cell-cell interfaces, many of which capture membrane and molecular dynamics but exclude hydrodynamics, or exploit equilibrium statistical physics and therefore omit dynamics, studying the influence of active forces requires a full fluid dynamics model. We have developed a computational fluid dynamics algorithm capable of simulating fluid-structure interactions with thermal fluctuations on seconds- and microns-scales [22]. Here, we use this to simulate two opposing membranes, variously including thermal fluctuations, active forces, and membrane permeability. We find that the thermal fluctuations are insufficient to overcome the thin-layer effect for a range of assumptions about molecular sizes. Active forces are sufficient to drive proximity. The thin-layer effect has the consequence of introducing two timescales (milliseconds and microseconds) in response to the two length scales inherent in the system. We find that membrane hydraulic permeability overcomes the thin-layer effects, but only for values larger than previous physiological estimates.

## Results

### Computational fluid dynamics simulation of the thin layer between cells

Receptor-ligand contact for the TCR occurs around 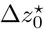 13 nm. Other parts of the membranes are separated by a distance Δ*z*_∞_, where estimates range from 22 nm to 150 nm [7, 23–29] for ectodomains of signaling molecules like CD45, non-specific binding pairs like LFA-ICAM and cadherins, and the glycocalyx. At the same time, cells themselves are *r*_cell_ ∼ 2 *μ*m for the smallest T cells [3]. To explore the consequences of the thin layer geometry, plus the incompressibility of fluid, we are required to simulate a 3D system with a resolution of receptor-ligand size 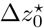 in a domain larger than the cell, which has radius *r*_cell_. (The analogous system in 2D would be insufficient since the opportunity for evacuating from the interface is fundamentally dependent on dimensionality of the boundary.)

We use an implementation of the stochastic immersed boundary framework to overcome these challenges, allowing us to simulate with parameters within the order-of-magnitude of experimentally estimated values, shown in Table 1. In this model, the two cell surfaces are represented by elastic disks, as shown in Fig. 1B, subject to bending resistance and approximate inextensibility. These disks are held by boundary tension *σ*_0_ in their plane, and separated by approximately inextensible nonspecific molecules of size Δ*z*_∞_, which we refer to as the far-field separation. These non-specific adhesions are absent from a region of radius *r*_free_ around the center of the disk, which we identify as the site of the receptor. The surface density of nonspecific adhesions is 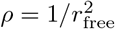. We assume both intracellular and extracellular fluids are Newtonian with viscosity of water, *η* = 10^−3^ Pa s. At the small length scales in our simulation, of ∼ nm, the viscosity of the cytosol can be one or two orders of magnitude larger [30], and at large length scales in our simulation, the viscosity is even larger. Thus, all times we report are underestimates, and dynamics at more realistic viscosity and cell separation are expected to be slower. Due to the linear nature of the fluid dynamics equations we use, all times scale linearly with viscosity.

**Table 1.** Model parameters

| Symbol | Name | Literature estimate<br>& source | Value<br>used here |
| --- | --- | --- | --- |
| $r_{\text{cell}}$ | Cell radius | $2 - 5 \mu\text{m}$ [3] | $1 \mu\text{m}$ |
| $r_{\text{free}}$ | LSM-free radius | $\sim 100 \text{ nm}$ [6] | $80 - 300 \text{ nm}$ |
| $\Delta z_\infty$ | Far-field separation, i.e., LSM height | $22 - 150 \text{ nm}$ [7, 23–29] | $30 - 120 \text{ nm}$ |
| $\Delta z_0^*$ | Critical separation for binding | $13 \text{ nm}$ [31] | $20 \text{ nm}$ |
| $\eta$ | Viscosity | $10^{-3} - 10^{-1} \text{ Pa s}$ [30] | $10^{-3} \text{ Pa s}$ |
| $B$ | Membrane bending modulus | $50 \text{ pN nm}$ [32] | $50 \text{ pN nm}$ |
| $\sigma_0$ | Membrane (boundary) tension | $0 - 100 \text{ pN}/\mu\text{m}$ [32, 33] | $0 - 100 \text{ pN}/\mu\text{m}$ |
| $F$ | Active force | $1 - 100 \text{ pN}$ [17, 18] | $1 - 20 \text{ pN}$ |
| $a$ | Force radius | $10 - 100 \text{ nm}$ [17, 34] | $10 \text{ nm}$ |
| $\psi$ | Membrane permeability | $10^{-2} - 10^1 \text{ nm}/\text{sPa}$ [19, 30] | $0 - 10^4 \text{ nm}/\text{sPa}$ |

### Thermal fluctuations are modulated by hydrodynamic dampening

As a control, we simulate a single membrane with thermal undulations, being held in place by adhesion molecules attached to fixed points in the fluid, as if it were attached to a “ghost” membrane. This simulation could be identified, for example, with a situation in which a cell is adhered to a highly permeable surface like a sparse network of extracellular matrix [35, 36] that provides minimal hydrodynamic confinement. A snapshot top view is shown in Fig. 2B. We find that the position of the receptor fluctuates as a Gaussian with standard deviation *σ* = 3.12 nm and an autocorrelation well-described by a single exponential decay with timescale *τ* = 1. 05 × 10^−6^s.

**Fig 2.**
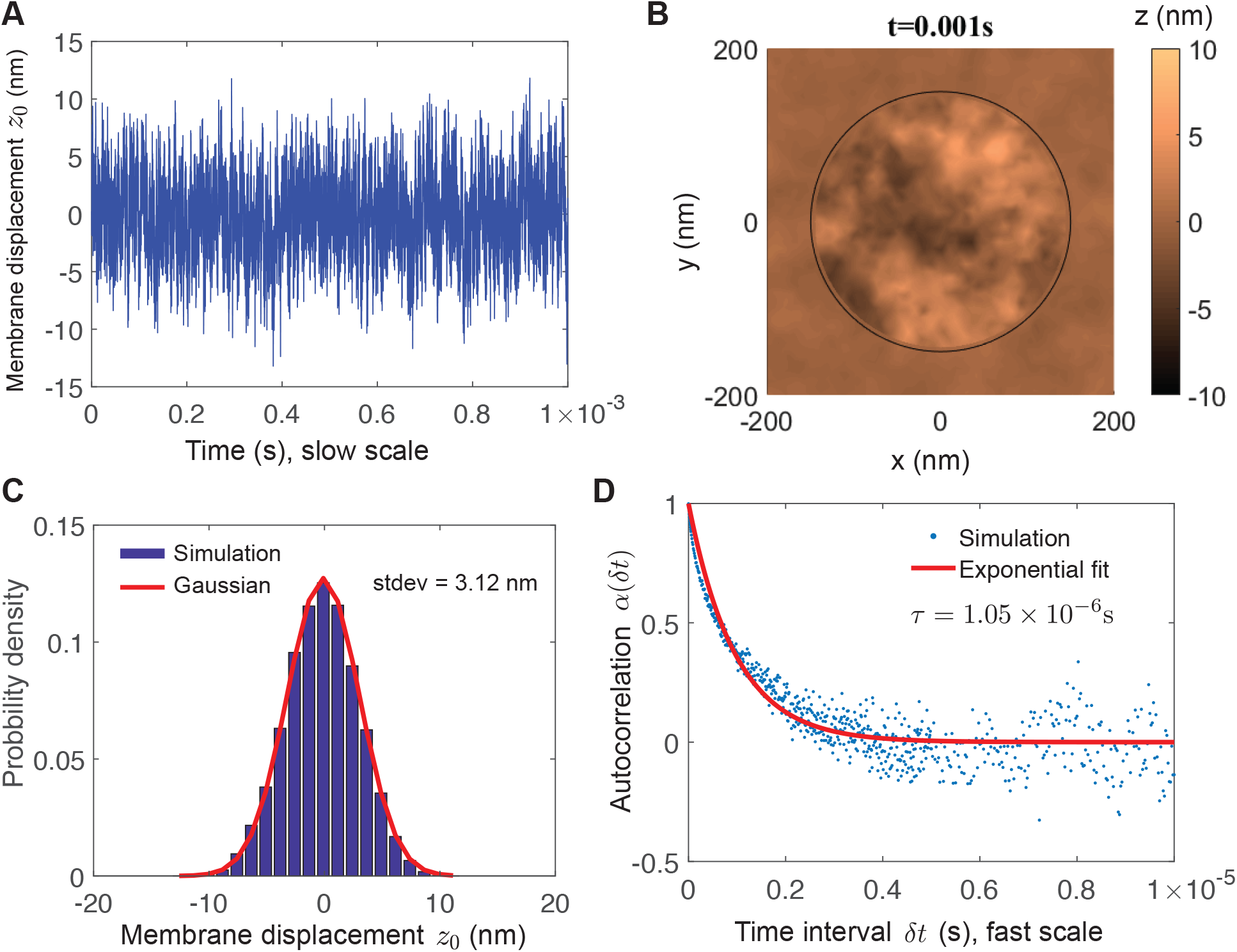
Thermal undulations of a single membrane. (A) Time series of membrane displacement at receptor coordinate *z*_0_(*t*). (B) Snapshot of membrane shape. For clarity the full simulation domain, extending *r*_cell_ = 1 *μ*m is not shown. (C) Stationary probability of membrane displacement at receptor coordinate follows a Gaussian distribution with zero mean and standard deviation 3. 12 nm. (D) Autocorrelation α(*δt*) of membrane displacement at receptor coordinate is well-approximated by a single exponential decay, indicating a simple stochastic process, with timescale *τ* = 1. 05 × 10^−6^s. Parameters used in this simulation are *r*_free_ = 150 nm, *σ*_0_ = 100 pN*/μ*m

We next simulate the interface with two membranes, as shown in Fig. 3D. The membranes are held at Δ*z*_∞_ = 60 nm outside the free radius. We run simulations for 1 s. We observe a stationary probability with mean separation 〈Δz〉 = 70. 0 nm. This blistering by 10 nm is due to an entropic repulsive pressure arising from thermal fluctuations [37, 38] and is not observed in simulations where thermal fluctuations are removed (Fig. 5B).

**Fig 3.**
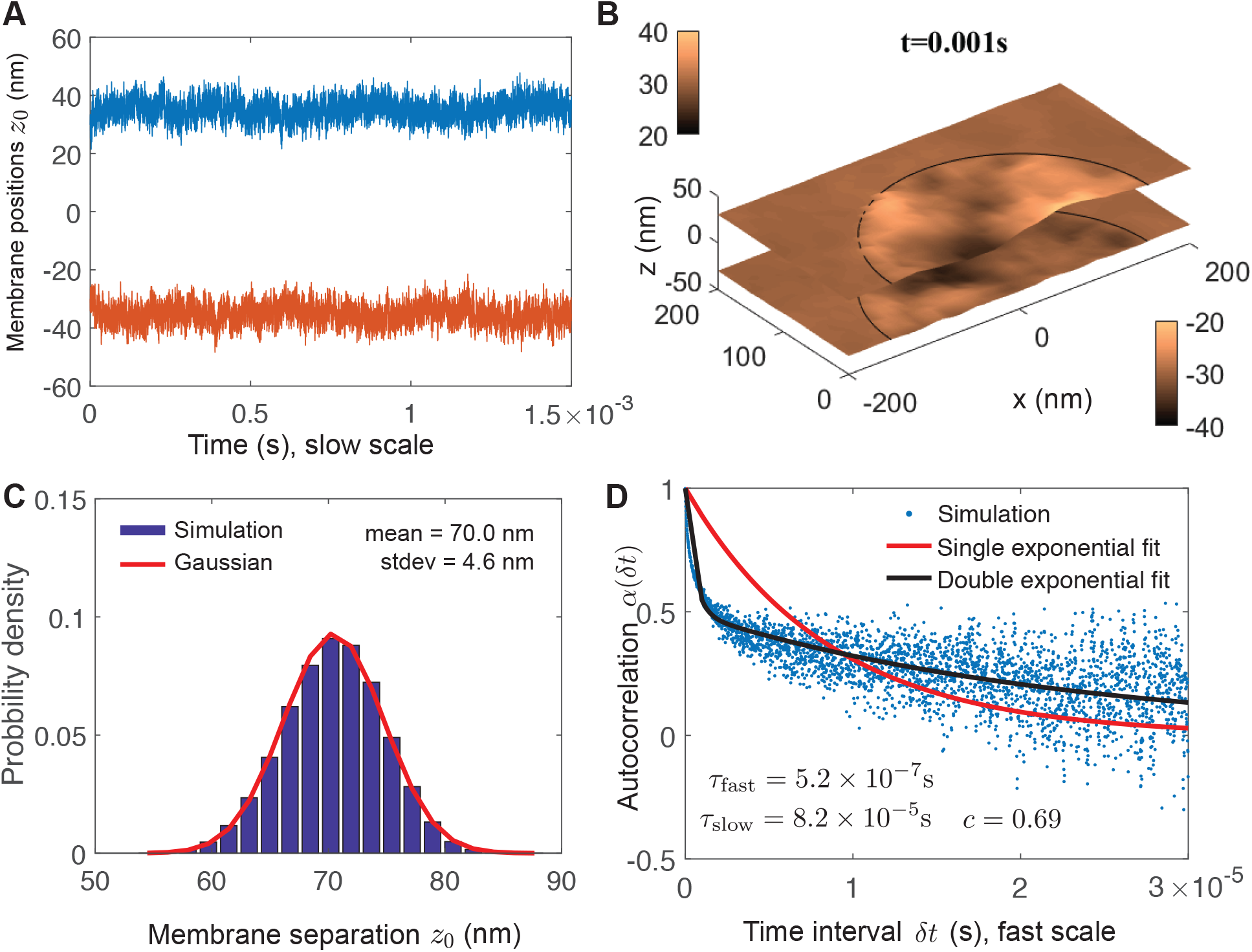
Thermal undulations of a cell-cell interface show thin-layer effect. Two membranes are held apart by Δ*z*_∞_ = 60 nm. (A) Time series of membrane positions at receptor coordinate. (B) Snapshot of membrane shapes. (C) Stationary probability of membrane separation at receptor coordinate follows a Gaussian distribution with mean 70 nm (larger than the far-field separation) and standard deviation 4. 6 nm. (D) Autocorrelation α(*δt*) of membrane separation does not fit a single exponential, but rather exhibits two timescales of decay, *τ*_fast_ = 5.2 × 10^−7^ s and *τ*_slow_ = 8.2 × 10^−5^s, where a fraction *c* = 0.69 of the composite process is attributed to the slow process. Note different time axis in (D) compared to Fig. 2D. Parameters used in this simulation are *r*_cell_ = 1 *μ*m, *r*_free_ = 150 nm, *σ*_0_ = 100 pN*/μ*m.

We observe a relatively small change in the amplitude of fluctuation compared to the single-membrane case, from 3.2 nm to 4.6 nm. The autocorrelation of Δ*z*_0_ does not fit a single exponential, but rather fits a two-timescale decay (black curve, Eq. 16) with a fast timescale *τ*_fast_ = 5.3 × 10^−7^ s comparable to the single-membrane autocorrelation above, but also a slow timescale *τ*_slow_ = 8.2 × 10^−5^ s. The double exponential equation we use to fit the autocorrelation is not a perfect fit, reflecting the inherent complexity of this process and the need for such computational modeling. This finding is in agreement with previous experimental work [11] showing that spatial amplitudes are not changed significantly, but fluctuation timescales are significantly altered by confinement.

### The timescales of thermal fluctuations are insufficient for close contact due to hydrodynamic dampening

Since the rate of receptor triggering is determined by the timescale of close contact, we next want to use the fluid dynamics simulations to estimate the mean first-passage time (MFPT) to close contact. Since these simulations include the target ligand only implicitly, we can infer the mean time to close contact for several values of 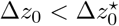. For the simulations with Δ*z*_∞_ = 60 nm, we ran simulations that for 1 second. For 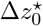 = 13 nm, close contact 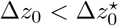 was not observed, suggesting it is a rare event in the sense that it occurs on a timescale much larger than the fluctuation timescale.

To overcome this computational challenge of observing such rare close contacts, we develop an approximation based on Ornstein-Ulhenbeck (OU) processes [39], and then use the Weighted Ensemble [40, 41] computational method to find the mean first time to a particular state of the system, here defined as the first time for the membranes to be within a distance of 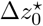 of each other. Full details are in Methods and S1 Appendix.

Because single-membrane dynamics exhibit a single timescale, the simplest model to explain both the stationary distribution of membrane distances and the autocorrelation function is a simple OU process, described by Eq. 13 in Methods, with parameters *σ* = 3. 12 nm and *τ* = 1. 05 × 10^−6^ s. At these parameters, we predict MFPTs shown in Fig. 4. For the single membrane case, an analytical approximation exists for the single-component OU [42], solid black line in Fig. 4, allowing us to confirm our computational method (further validation is provided in S1 Appendix)

**Fig 4.**
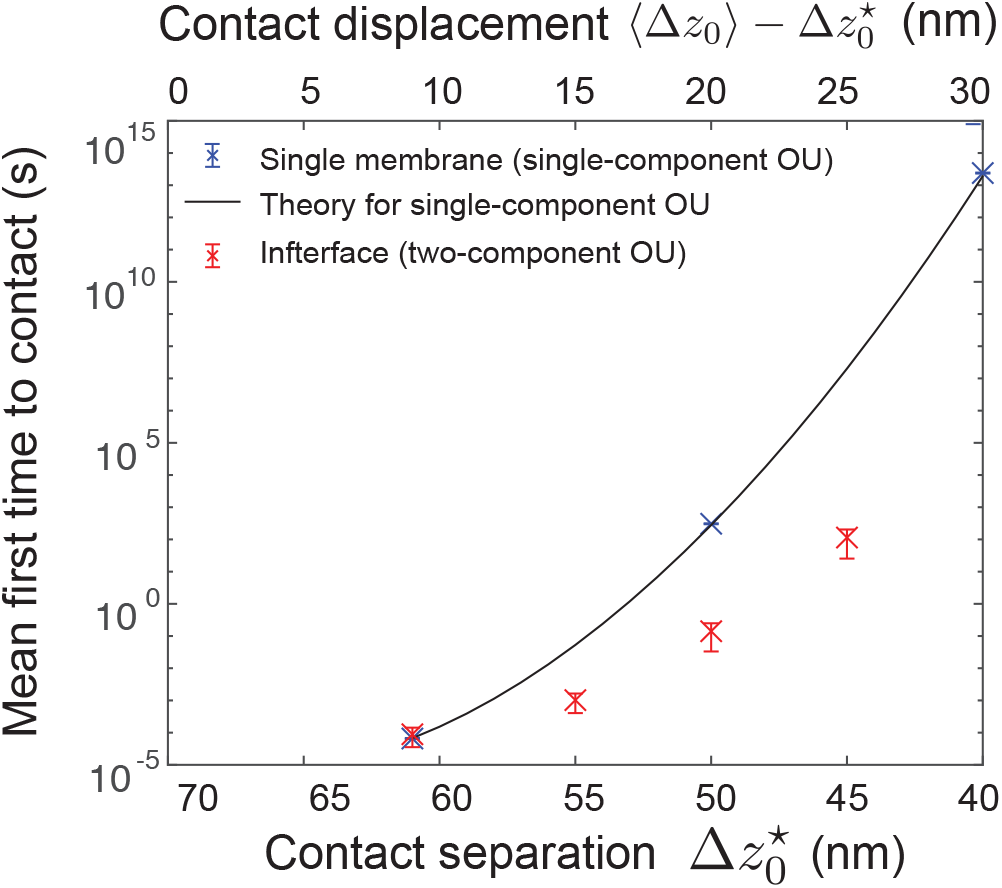
Approximation of mean first-passage time to close contact. (For a single membrane with parameters from Fig. 2C,D, the MFPTs (blue) agree with theoretical results from [42]. For two membranes with parameters from Fig. 3C,D, the MFPTs (red) increase super-exponentially.

For the interface, we find that membranes will displace by 20 nm (i.e., the separation distance deviated from its mean of 70 nm down to 50 nm) in approximately 1 second. The time until a displacement larger than this grows super-exponentially: for a displacement of 25 nm (i.e., down to separation 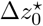 = 45 nm), it takes ∼ 100s.

In Fig. 4, the interface case apparently has a larger (i.e., slower) MFPT for the single membrane. However, we note that these numbers are not directly comparable. The single-component OU describes the position of a single membrane, which has standard deviation *σ*_1_ = 3. 1 nm, while the two-component OU describes the distance between two membranes, which has a standard deviation *σ*_2_ = 4. 6 nm. A more direct comparison would be a hypothetical simulation in which two “single” membranes were held at a distance of 70 nm, but did not interact via fluid therefore would fluctuate independently. In such a case, the separation between these membranes would be 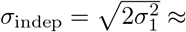 4.4 nm, approximately the same as the interface standard deviation.

### Active forces from F-actin filopodia-like protrusions are significantly hampered by interface but still sufficient for rapid close contact

Cells, including the T cell, continuously extend active processes driven by F-actin like filopodia and microvilli [34, 43] that facilitate receptor binding [29, 44]. To explore the effect of hydrodynamics on active processes at an interface, we simulate a force *F* at the receptor site, spread over a disk of radius *a* = 10 nm 1B. In Fig. 5, we find that a force of *F* = 20 pN is sufficient to drive close-contact from a far-field separation of Δ*z*_∞_ = 50 nm for both single membranes and interfaces.

We perform deterministic simulations with thermal forces omitted (black curves). The dynamics are quantitatively similar, and the simulations are much less computationally taxing. For this reason, for the remainder of this section we perform simulations without thermal fluctuations. Note in Fig. 5 the stochastic and deterministic simulations approach equilibrium on approximately the same slow timescale, but the equilibrium separation is larger when thermal forces are included due to the entropic repulsion discussed above.

**Fig 5.**
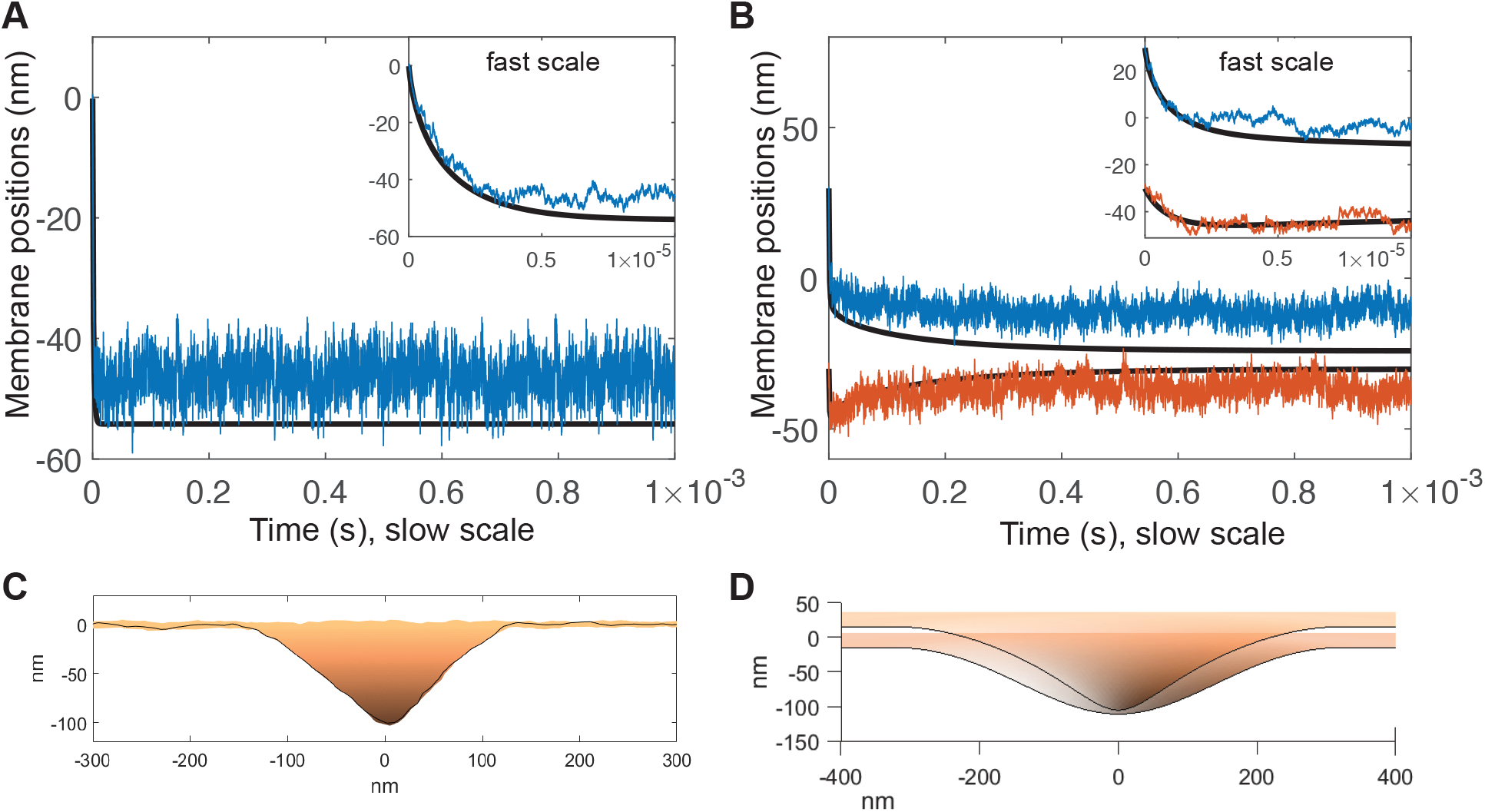
Active forces driving membrane proximity. (A) Active force of *F* = 20 pN applied to single membrane. Simulations including thermal undulations (blue) compared to purely deterministic simulations without thermal undulations. Inset shows fast timescale of mechanical equilibration. (B) Active force of *F* = 20 pN applied to top membrane at a cell-cell interface held apart Δ*z*_∞_ = 50 nm. After rapid initial phase (inset), equilibrium separation is not reached until ∼ 10^−3^s. Parameters used in this simulation are *r*_cell_ = 1 *μ*m, *r*_free_ = 150 nm, *σ*_0_ = 100 pN*/μ*m. (C) Snapshot of equilibrium from single-membrane simulation with thermal fluctuations. (D) Snapshot of intermediate configuration at *t* = 10^−4^s from interface simulation without thermal fluctuations.

The shape of the protrusion is shown in Fig. 5C,D. Membrane profiles are reminiscent of micrographs of microvilli in T cells (see, e.g., [29] Fig. 3G): The edges are rounded due to membrane bending resistance, and closest contact is at the tip, with cell separation distance gradually tapering off.

To isolate the influence of the thin-layer effect, we perform identical simulations with and without a second membrane, for various active forces, in Fig. 6. For a single membrane, the distance approaches a new equilibrium rapidly, ∼ 10^−5^s. For an interface, there is an initial rapid movement of the top membrane, i.e, the driven membrane (blue curve in B) ∼ 10^−5^s, however this is accompanied by a rapid depression of the bottom membrane, i.e., the passive (red curve). Then, on a slower timescale ∼ 10^−3^s, the passive membrane returns. We attribute the rapid depression to the incompressibility of the extracellular fluid, and the slow timescale to the thin-layer timescale identified above, as the excess fluid must drain from the interface.

**Fig 6.**
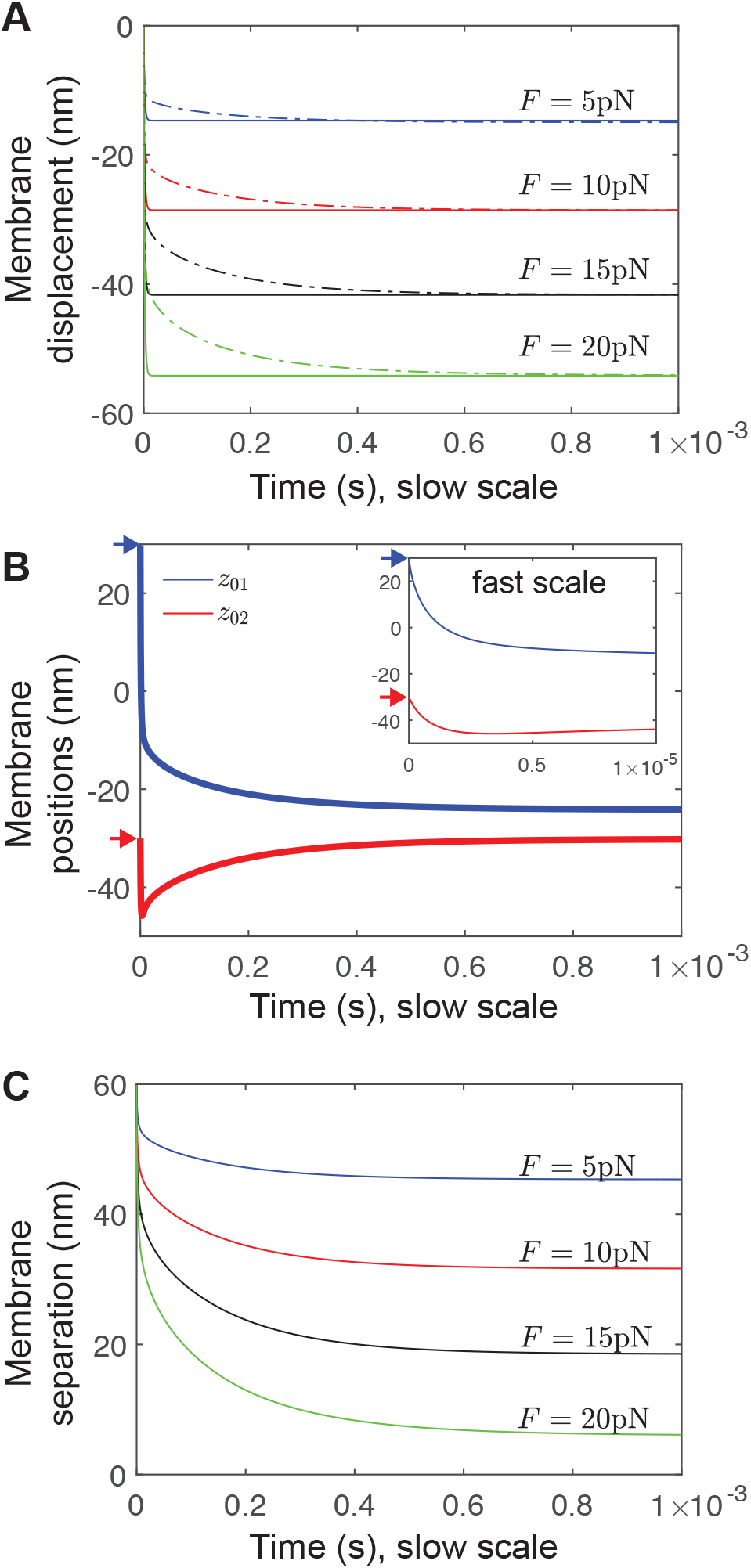
Active forces at a cell-cell interface exhibit a slow timescale of equilibration due to thin-layer effect. (A) Membrane displacement for various forces for a single membrane (solid curves) and at an interface (dashed curves). (B) Membrane positions for *F* = 10 pN. The top membrane (blue) moves in an manner initially similar to the single-membrane case (inset), while the bottom membrane (red) is pushed away by hydrodynamic interaction. Then, on the slow timescale, the bottom membrane moves back up towards its equilibrium. Arrows indicate initial positions, to highlight the rapid initial movement otherwise difficult to see. (C) Membrane separation (which, in contrast to membrane displacement in (A), includes the slow return of the bottom membrane) for various forces.

### Influence of membrane tension and density of surface molecules

The plasma membrane is under tension, maintained by hydrostatic pressure and regulation of exocytosis, endocytosis and membrane ruffles [32, 45] and is in the range of 3 *−* 300 pN*/ μ*m [32, 33] and it spatially nonuniform [46]. We apply membrane tension in our simulation as a boundary surface tension with magnitude *σ*_0_. We find that higher surface tension necessitates more force for the equivalent equilibrium displacement, as shown in Fig. 7A. This demonstrates that the system is above the critical length scale below which surface tension is insignificant compared to membrane bending [17, 47]. Note that these these data show the equilibrium position in response to a constant force, therefore there is no effect of fluid dynamics, and thus no thin-layer effect.

**Fig 7.**
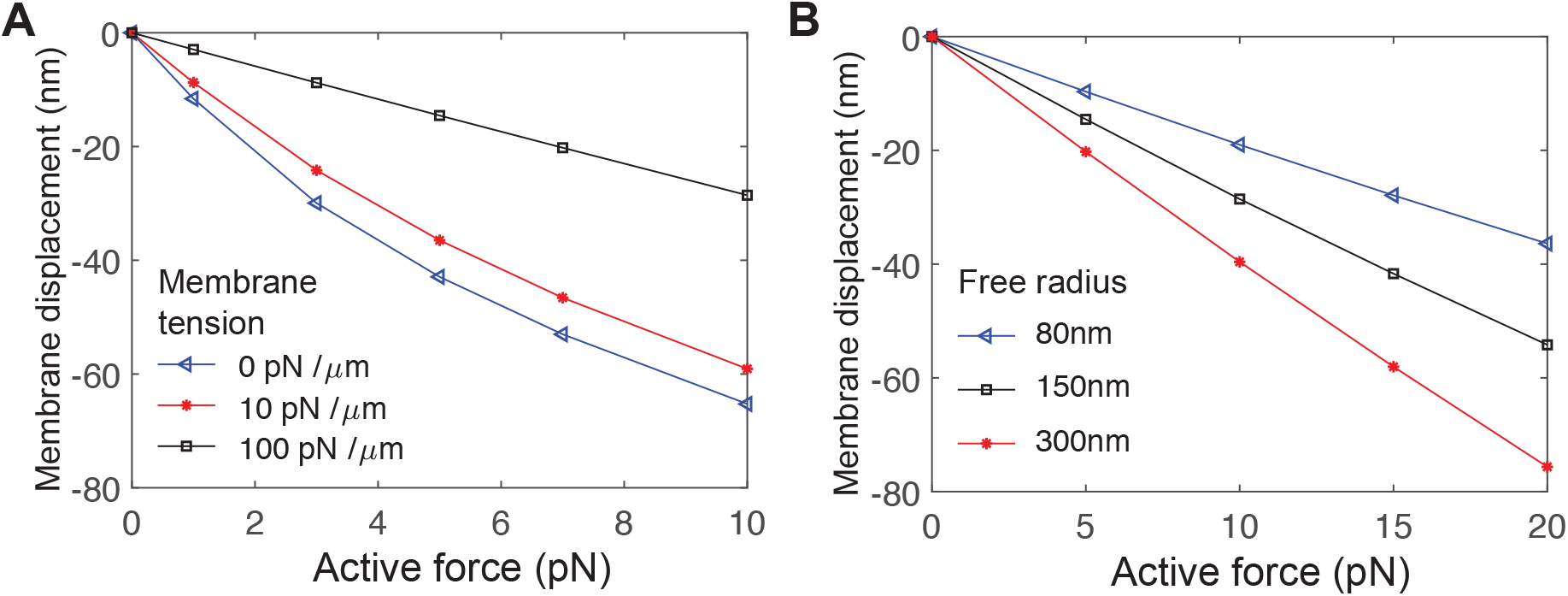
Equilibrium membrane displacement as a function of active force. (A) For various surface densities of non-specific adhesion molecules *ρ*, which relates to the free radius near the receptor *r*_free_ by 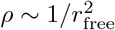 (B) For various values of membrane tension *σ*_0_, applied at the boundary of both cells. Note since these data are at equilibrium, there is no thin-layer effect, and membrane separation is far-field separation minus membrane displacement.

The results we report are sensitive to the properties of the large surface molecule, such as its surface density *ρ* or equivalently its intermolecular spacing *r*_free_. We previously estimated that close contact occurs in depletion zones with *r*_free_ ∼ 100 nm [6]. In Fig. 7B, we explore the equilibrium separation as a function of force for various surface densities, *r*_free_ = 80 nm(*ρ* = 1.6 × 10^−4^ nm^−2^), *r*_free_ = 150 nm(*ρ* = 4.4 × 10^−5^ nm^−2^), *r*_free_ = 300 nm(*ρ* = 1.1 × 10^−5^ nm^−2^). As expected, higher density of surface molecules reduces the equilibrium displacement.

### Optimal distance away from the target cell for active extension

The results we report are also sensitive to the molecular size Δ*z*_∞_ of the large surface molecule. The large range of estimates for Δ*z*_∞_ arises from the uncertainty about which molecules dominate the process of keeping the membranes apart. Molecules like CD45 may sterically maintain membrane separation by as little as 22 nm. Non-specific adhesion molecules like LFA-ICAM and cadherins are estimated to span a range from 28 nm [23] to 43 nm [24, 25]. Estimates for the thickness of the glycocalyx range from 40 – 50 nm [26, 27] to 150 nm [28, 29]. So, we explore receptor proximity driven by active forces, varying the far-field separation Δ*z*_∞_ from 30 nm to 120 nm in Fig. 8.

**Fig 8.**
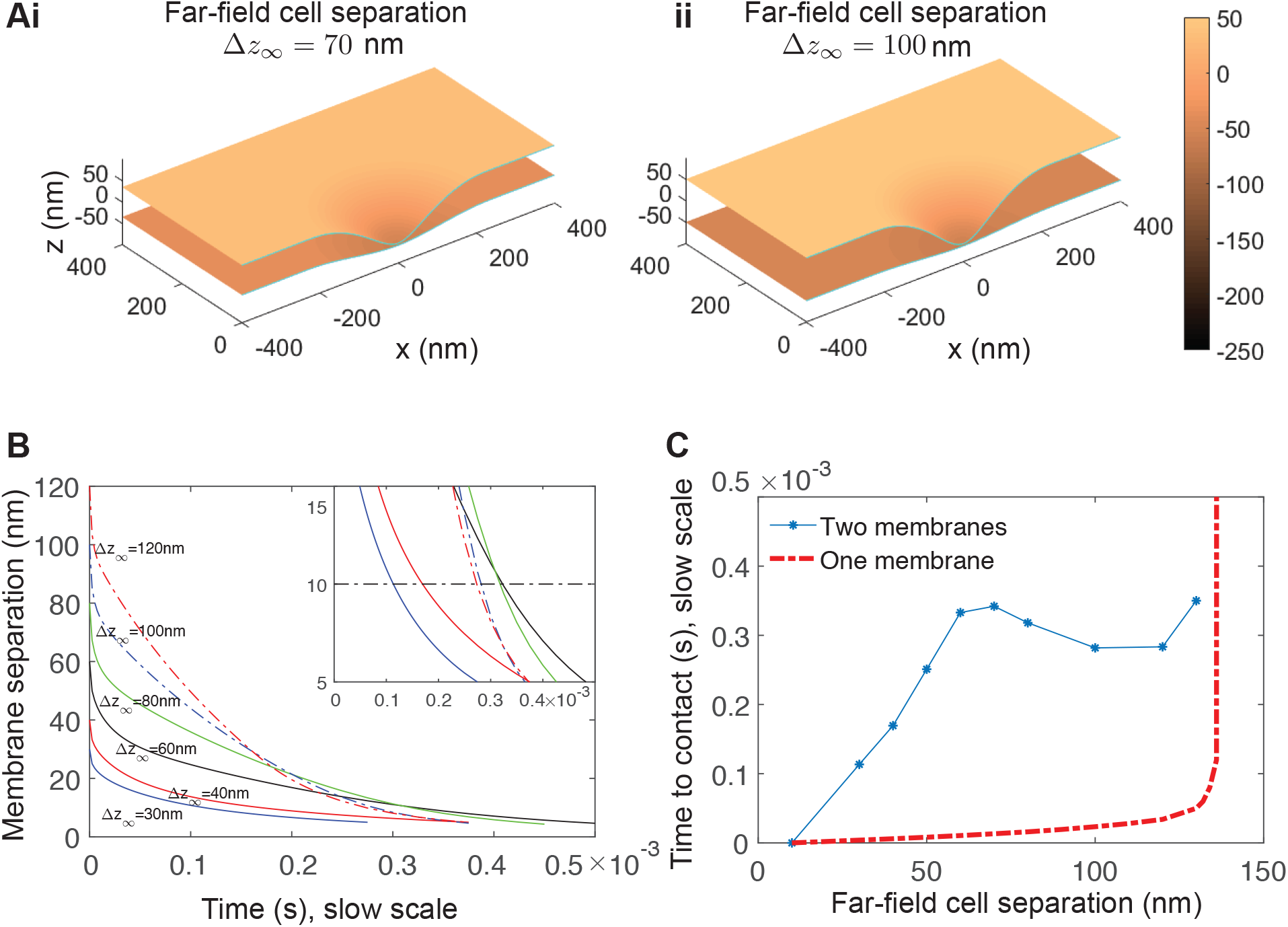
Active force of 10 pN for various initial cell separation distances demonstrates an optimal initial distance. (A) Snapshot of simulations with far-field separation Δ*z*_∞_ = 70 nm and Δ*z*_∞_ = 100 nm. Parameters used for both are *r*_cell_ = 1 *μ*m (for clarity the full simulation domain is not show), *r*_free_ = 200 nm, *σ*_0_ = 0. At these parameters, this force is sufficient to drive Δ*z* < 10 nm. (B) Membrane separation for various far-field separation Δ*z*_∞_ = 70 nm. Inset shows non-monotonic behavior where the time series cross. (C) Time until Δ*z* < 10 nm versus far-field separation (blue asterisks). Simulations for a single membrane are shown (red, dashed) for comparison. The single-membrane time diverges around 140 nm since this force induces an equilibrium deformation of that magnitude (Fig. 7).

As the starting distance is increased, the time before proximity 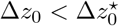 increases, shown in Fig. 7B. However, we find that above a critical far-field separation Δ*z*_∞_ ≈ 80 nm, membrane deformation speed increases, even though force is kept constant at *F* = 10 pN.

The effect is modest but sufficient so that, over a large range of far-field separations ∼ 60 – 130 nm, the time to contact does not increase for increasing separation, Fig. 7C. The optimal distance is approximately independent of force magnitude (not shown). Heuristically, this plateau arises because of a significant thin-layer effect dominates motion. Since this effect depends sensitively on the thickness of the thin layer, increasing the thickness reduces the effect, and the active protrusion can push more easily through the free fluid. On the other hand, although speed increases, the distance to the target cell also increases. Thus there is an optimal “attack distance” from which to extend a protrusion.

### Membrane permeability value for which thermal and active proximity is accelerated

If a membrane were perfectly water permeable *ψ* = ∞ there would be no thin-layer effect. Biological membranes are sufficiently permeable that, in fast motile cells, fluid velocity appears stationary in the lab frame of reference [19]. Therefore, it is a priori reasonable to expect that there is a magnitude of permeability above which the slow-timescale behavior of the interface is removed, leaving only fast dynamics. We repeat the active force simulations at *F* = 20 pN, and explore permeabilities at each order of magnitude, in Fig. 9. We find that the first significant deviation from impermeability (*ψ* = 0) occurs at *ψ* ∼ 10^2^ nm/s Pa (red dotted). By *ψ* = 10^4^ nm/s Pa, the top membrane time series is comparable to the single-membrane case (Fig 6B), i.e., very little thin-layer effect is observed. The transition to fast-only dynamics occurs through a reduction in timescale, and only a weak reduction in amplitude (green curve is compressed horizontally).

**Fig 9.**
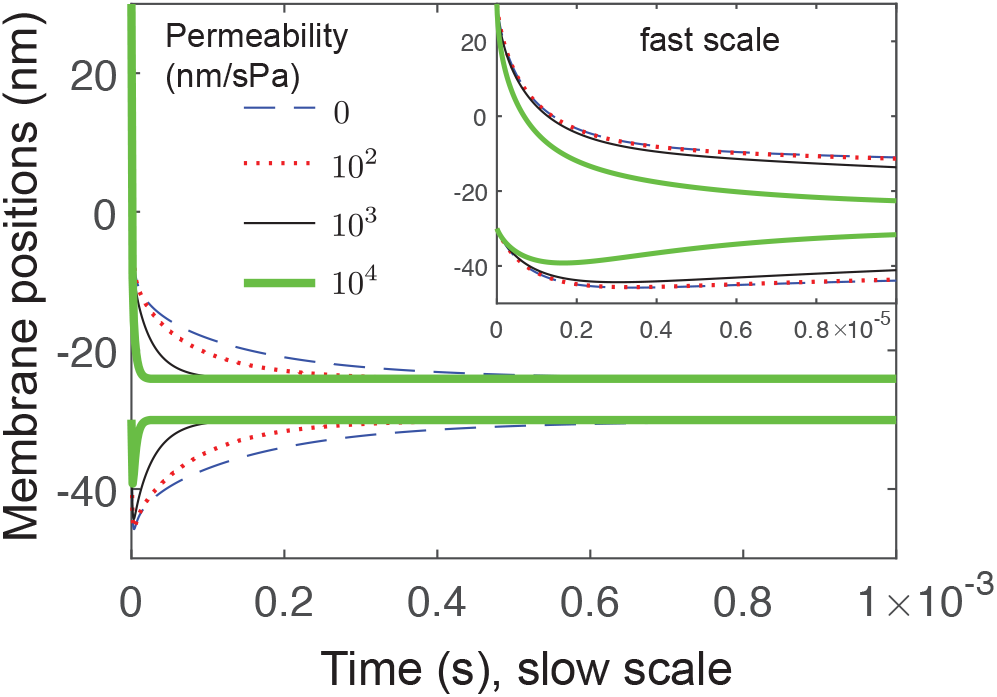
Influence of membrane permeability on thin-layer effect. Membrane positions are shown for *F* = 10 pN. For zero permeability (blue dashed), there is significant thin-layer effect. Significant changes to the time series are first seen for permeability *ψ* = 10^2^ nm/s Pa (red dotted). By permeability *ψ* = 10^4^ nm/s Pa, the top membrane time series is comparable to the single-membrane case (Fig 6B).

This is orders of magnitude larger than the permeability estimates 10^−2^ nm/s Pa[30]. The largest indirect estimates from motile epithelial keratocytes gives ∼ 10^1^ nm*/* s Pa [19]. So, taken together, our results suggest the thin-layer effect cannot be abrogated by physiological levels of permeability.

## Discussion

Fluid dynamics plays a role in cellular processes like the swimming of eukaryotes and bacteria, ciliary beating [48], and cell blebbing [49, 50], but also in less obvious examples like nuclear shape [51], organelle positioning [52, 53], some surface crawling of both eukaryotes [54] and bacteria [55], and ultra-fast endocytosis in neurons [56]. This work studies fluid dynamic effects in the context of transient cell-cell contact by immune cells [11, 13]. We find that thermal fluctuations alone are reduced in speed, but not amplitude, in agreement with previous experimental work [11], that active forces are sufficiently fast but still significantly slowed, and that physiological levels of membrane permeability do not significantly change this.

Active protrusive forces like filopodia and microvilli are abundant in cells including naive, resting and activated T cells [29, 44]. However, in vitro reconstitutions in which two cell-sized lipid vesicles are brought into contact [10] do not have active protrusion. Our simulations without active protrusion (Fig. 3) predicts significant delay before the first reports of receptor-ligand contact. Interestingly, in vitro reconstitution take approximately 16 minutes before signs of molecular binding [10]. Our work indicates a source of this delay is the long timescale of fluctuations due to the thin layer: Hydrodynamics in the interface between vesicles is slow, and in the total absence of active protrusion, receptor proximity must rely on thermal fluctuations hampered by the thin-layer effect.

For cells with active protrusions, our results suggest that the thin-layer effect can be readily overcome by the typical forces of filopodia [17, 18]. In our model, a constant force is maintained by the protrusive machinery as the membrane is driven outward, implicitly assuming that hydrodynamic drag is the rate-limiting process. Protrusions driven by polymerization of F-actin must assemble monomers at the leading edge, a process which can drive protrusions at 200 – 300 nm/s [17,19], speeds that could produce close-contact from a distance of Δ*z*_∞_ = 50 nm in ∼ 10^−1^1s. The hydrodynamic-limited case we explore here produces close-contact at this distance in ∼ 10^−3^s, suggesting that hydrodynamics is not rate-limiting. However, we note that our simulations assumed both the cytosolic and extracellular viscosities are that of water, η = 10^−3^ Pa s. This is a conservative estimate compared to established measurements of cytosolic viscosity that are one or two orders of magnitude larger [30]. Repeating our simulations with a change in viscosity would linearly scale all times, so a tenfold increase in viscosity would slow contact by tenfold. In this case, it is possible that hydrodynamics becomes limiting.

The slow timescale of hydrodynamic relaxation could explain the appearance of secretory clefts [57], long-lived blisters of extracellular fluid that are hypothesized to be particularly important for cytotoxic T cell function, since they ensure cytolytic granules secreted by the T cell are concentrated near the target cell [3, 58]. These blisters may arise and persist out-of-equilibrium due to the long-timescale of fluid evacuation through the tight cell-cell contact regions. This would provide an example of a cell-biological structure arising as a consequence of simple fluid dynamics, upon which regulation occurs by structures like the microtubule organizing center [58, 59].

A major opportunity provided by computational fluid dynamics studies, rather than, e.g., analytical approaches, is the study of more realistic geometry with more molecular participants, such as the F-actin cortex and its adhesion with the plasma membrane [50,60]. We expect that modulating cortex-membrane adhesion would allow us to simulate active protrusions that, at high adhesion, behave more like narrow, finger-like filopodia [17], while at low adhesion behave more like microvilli and invadopodia [29], which are rounder and wider [44].

## Methods

### Numerical implementation

We use the Stochastic Immersed Boundary Method, an extension if the Immersed Boundary Method [61] developed by Atzberger and coworkers [62–64]. The fluid has velocity field **u** parameterized by Eulerian coordinate **x** ∈ . The immersed structure has configuration described by **X** and is parametrized by *s* ∈  in the membrane domain . The equations of motion are 
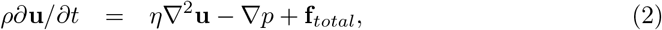

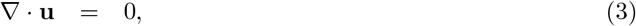

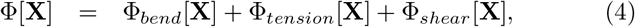

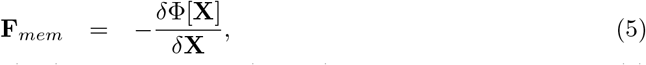

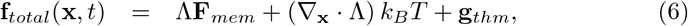

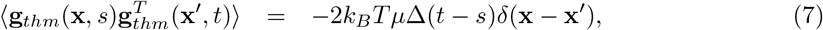

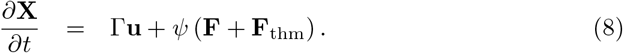

The first term in Eq. 2 is the inertial term, where *ρ* is the fluid mass density, which must be included to accommodate thermal fluctuations (even though simple scaling put the system in the low-Reynolds regime [22, 62]). The pressure *p* is imposed by the incompressiblity condition in Eq. 3. Eq. 7 describes the stochastic driving fields, which are chose to obey the fluctuation-dissipation principle [63, 64].

The force from the membrane, Eq. 5, is computed using a variational approach from the membrane energy. We describe the Helfrich energy functional [65] with bending energy 
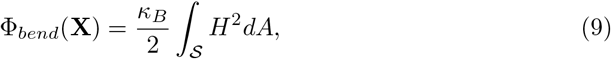
where *κ_B_* is bending rigidity and *H* is mean curvature of the membrane surface. In addition to the bending energy, we consider a membrane that resists area changes by a surface tension energy 
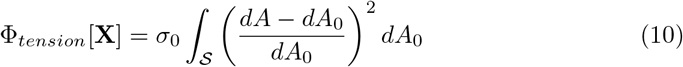
where *σ*_0_ is the surface tension constant. Finally, the membrane resists shear in order to maintain numerical stability [64].

Eq. 8 describes the motion of the membrane, which follows the fluid velocity *u* but with a pressure-driven difference due to permeability, with coefficient *ψ*.

The fluid and structure are coupled by 
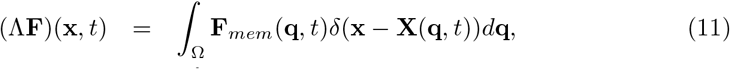

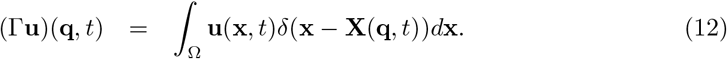

Full description of energy terms and details of numerical implementation are in the S1 Appendix.

### Estimation of mean first-passage time

In the absence of deterministic forces, the receptor-site membrane distance Δ*z* follows a stochastic trajectory. We find that, in the case of a single membrane, it is well-approximated by an Ornstein-Ulenbeck process [39] in the new variable *Z* = Δ*z* – 〈Δ*z*〉, 
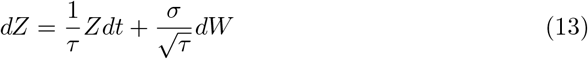
where *W* is a Weiner process. This has stationary distribution 
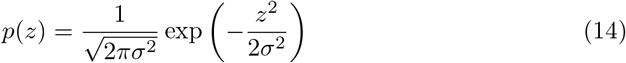
and autocorrelation function 
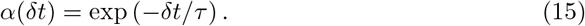

For the case of an interface, we find that the receptor-site membrane-membrane distance Δ*z* follows a stochastic trajectory with autocorrelation that is well-approximated by 
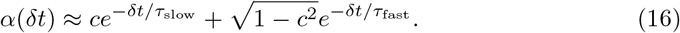

This is the autocorrelation function of a two-component OU process 
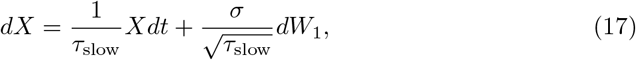

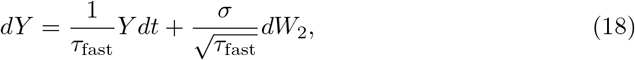

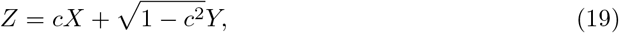
where, without loss of generality, we assume the two hidden components *X* and *Y* have the same variance *σ*^2^ (since any difference can be absorbed into *c*), and we define the fraction of the process attributed to the slow timescale *c* and fast timescale 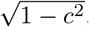 again without loss of generality, as a convenient way of fitting the variance.

We refer to Eq. 13 as the one-component OU process to describe a single membrane, and Eqs. 17-19 as the two-component OU process to describe the interface. For both of 324 these, we present methods for determining MFPTs in the S1 Appendix.

## Supporting information 32

**S1 Appendix. Supplemental methods.** Description of computational fluid dynamics method and implementation. Description of weighted ensemble method and implementation.

## Acknowledgments

This work was supported by NSF CAREER grant DMS 1454739 to JA, NSF grant DMS 1715455 to ELR, NSF grant DMS 1763272 and a grant from the Simons Foundation (594598, QN).

